# Disruption of riboflavin biosynthesis in mycobacteria establishes 5-amino-6-D-ribitylaminouracil (5-A-RU) as key precursor of MAIT cell agonists

**DOI:** 10.1101/2024.10.03.616430

**Authors:** Melissa D. Chengalroyen, Nurudeen Oketade, Aneta Worley, Megan Lucas, Luisa Nieto Ramirez, Mabule L. Raphela, Gwendolyn M. Swarbrick, Digby F. Warner, Deborah A. Lewinsohn, Carolina Mehaffy, Erin J. Adams, William Hildebrand, Karen Dobos, Valerie Mizrahi, David M. Lewinsohn

**Affiliations:** UCT Molecular Mycobacteriology Research, Institute of Infectious Disease and Molecular Medicine, Department of Pathology, University of Cape Town, Cape Town, South Africa; Department of Microbiology, Immunology and Pathology, Colorado State University, Colorado, United States of America; Oregon Health and Science University, Oregon, United States of America; Wellcome Centre for Infectious Disease Research in Africa, University of Cape Town; Department of Biochemistry and Molecular Biology, University of Chicago, Chicago, Illinois, United States of America; Department of Microbiology and Immunology, University of Oklahoma Health Sciences Centre, Oklahoma City, Oklahoma, United States of America; Portland VA Medical Center, Oregon, United States of America

## Abstract

Mucosal-associated invariant T (MAIT) cells exhibit an intrinsic ability to recognize and respond to microbial infections. The semi-invariant antigen recognition receptor of MAIT cells specifically detects the non-polymorphic antigen-presenting molecule, major histocompatibility complex class I-related protein 1 (MR1), which primarily binds riboflavin-derived metabolites of microbial origin. To further interrogate the dependence of these antigens on riboflavin biosynthesis in mycobacteria, we deleted individual genes in the riboflavin biosynthesis pathways in *Mycobacterium smegmatis* (Msm) and *Mycobacterium tuberculosis* (Mtb) and evaluated the impact thereof on MAIT cell activation. Blocking the early steps of the pathway by deletion of RibA2 or RibG profoundly reduced, but did not completely ablate, MAIT cell activation by Msm or Mtb, whereas deletion of RibC, which catalyzes the last step in the pathway, had no significant effect. Interestingly, deletion of RibH specifically enhanced MAIT cell recognition of Mtb whereas loss of lumazine synthase (RibH) activity had no impact on MAIT cell activation by Msm. MAIT cell activation by Msm was likewise unaffected by blocking the production of the MAIT cell antagonist, F_o_ (by inhibiting its conversion from the riboflavin pathway intermediate, 5-amino-6-D-ribitylaminouracil (5-A-RU), through the deletion of *fbiC)*. Together, these results confirm a central role for 5-AR-U in generating mycobacterial MR1 ligands and reveal similarities and differences between Msm and Mtb in terms of the impact of riboflavin pathway disruption on MAIT cell activation.

**Author summary:** Mucosal-associated invariant T (MAIT) cells are an abundant population of innate-like T-cells which respond to microbial infections. These specialized cells recognize the MR1 molecule, which presents microbial metabolites derived from riboflavin (vitamin B2) biosynthesis. These cells are enriched in the airways and in some cases reduced in the peripheral blood of tuberculosis (TB) infected individuals suggestive of a role in the early response to infection by *Mycobacterium tuberculosis*. In this study, we investigated the effect of deleting individual genes in the riboflavin biosynthesis pathway on MAIT cell activation by *Mycobacterium tuberculosis* or *Mycobacterium smegmatis*. Our findings revealed that disrupting early stages in the pathway profoundly reduced but did not eliminate MAIT cell activation by both mycobacterial species. However, blocking the penultimate step in the pathway, catalyzed by the enzyme, lumazine synthase, led specifically to increased MAIT cell recognition of *M. tuberculosis*. Our results confirm the pivotal role of the riboflavin pathway intermediate, 5-A-RU, in generating mycobacterial ligands that serve as MAIT cell agonists. By enhancing our understanding of how MAIT cells recognize mycobacterial infections, the results of this study could inform strategies for the development of vaccines and/or immunotherapies for TB.

## Introduction

MR1-restricted T cells (MR1Ts) are characterized by their dependence on the highly conserved molecule, MR1 [1]. A subset of MR1Ts, Mucosal Associated Invariant T cells (MAITs), have been characterized by their usage of a semi-invariant T cell receptor (TRAV1-2) [1, 2], their expression of the transcription factor PLZF, and expression of CD161 and CD26 [3]. While these cells are among the most abundant in the human circulation and in mucosal sites, their function was largely unknown until 2010 when they were found to recognize a variety of microbes, including the major human pathogen, *Mycobacterium tuberculosis* (Mtb) [4, 5]. A seminal discovery was made in 2012 when Kjer-Nielsen *et al*. [6] demonstrated that MR1 could present the metabolite, 5-(2-oxopropylideneamino)-6-D-ribitylaminouracil (5-OP-RU), derived from an intermediate in the riboflavin biosynthesis pathway of *Salmonella* sp., to MAIT cells. These authors argued that microbes producing riboflavin are uniquely poised to activate these cells. Microbially derived MR1 ligands and antigens have since been found to be more diverse than originally described [7]; others that have been discovered include photolumazines I, III and V, which were identified in *Mycobacterium smegmatis* (Msm) and are also derived from the riboflavin pathway [8, 9]. Furthermore, the source of MR1 ligands that are stimulatory for MAIT cells may extend beyond the riboflavin pathway as evidenced by our finding that the TRAV1-2-negative, TRAV12-2-expressing T-cell clone, D462-E4, recognizes ligands derived from *Streptococcus pyogenes*, a riboflavin auxotroph [10]. This suggests that MR1Ts can distinguish between microbes in a TCR-dependent manner.

The central role of the microbial riboflavin pathway in MAIT cell activation has been confirmed in different organisms. These include deletion mutants of *Lactococcus lactis* [7], *Salmonella* Typhimurium [11], *Escherichia coli* [12], *Streptococcus pneumoniae* [13]and *Aspergillus fumigatus* [14] rendered auxotrophic for riboflavin as well as natural variants of *S.* Typhimurium [15] and *S. pneumoniae* [13] with altered expression of the riboflavin pathway. Moreover, engineered overexpression of specific riboflavin pathway genes in Msm, Mtb or *M. bovis* BCG was shown to enhance MAIT cell activation, albeit differentially between the three mycobacterial species and to varying extents, depending on the gene targeted for over-expression [16]. Building on our prior work demonstrating the essentiality of riboflavin biosynthesis in Mtb and Msm and the ability of both organisms to transport and assimilate riboflavin [17], we set out to investigate the impact of riboflavin pathway disruption on MAIT cell activation by these mycobacteria. We generated mutants of Mtb and Msm auxotrophic for this cofactor by knocking out the biosynthesis pathway genes and analyzed the impact thereof on the expression of the pathway genes and proteins, and on MAIT cell activation. We show that blocking the early steps of the pathway by deletion of RibA2 or RibG profoundly reduced but did not completely eliminate MAIT cell activation whereas deletion of RibC, which catalyzes the last step in the pathway, had no discernable effect on MAIT cell activation by Msm or Mtb. Moreover, deletion of RibH specifically enhanced MAIT cell recognition of Mtb whereas loss of lumazine synthase activity had no significant impact on MAIT cell activation by Msm. MAIT cell activation by Msm was likewise unaffected by blocking the production of the MAIT cell antagonist, F_0_, derived from the riboflavin pathway intermediate, 5-A-RU by deletion of *fbiC*. Our results confirm a central role for 5-A-RU in the production of mycobacterial MR1 ligands and reveal similarities and differences between Msm and Mtb in terms of the impact of riboflavin pathway disruption on MAIT cell activation. These findings will guide future studies in the field of MR-1 immune responses towards Mtb.

## Results

### Construction of deletion mutants in the riboflavin pathway of Msm and Mtb

In a recent study [17], we characterized the riboflavin biosynthesis pathway in Mtb H37Rv and Msm mc^2^155 and confirmed that it comprises a bifunctional GTP cyclohydrolase II/3,4-DHBP synthase (RibA2); riboflavin deaminase/5-amino-6-(5-phosphoribosylamino) uracil reductase (RibG); an unidentified phosphatase; lumazine synthase (RibH), and riboflavin synthase (RibC) (**Fig 1A and 1B**). Riboflavin is the precursor of flavin mononucleotide (FMN) and flavin adenine dinucleotide (FAD), which are produced by the bifunctional enzyme, RibF. The biosynthetic pathway also leads to the production of the deazaflavins, F_0_ and F_420_, from the pathway intermediate, 5-A-RU, via the sequential action of the enzymes FbiC, FbiA and FbiB [18]. Together, the flavin and deazaflavin cofactors enable the activity of a myriad enzymes in mycobacteria, organisms which adopt ‘flavin-intensive’ lifestyles [19, 20].

**Fig 1.**
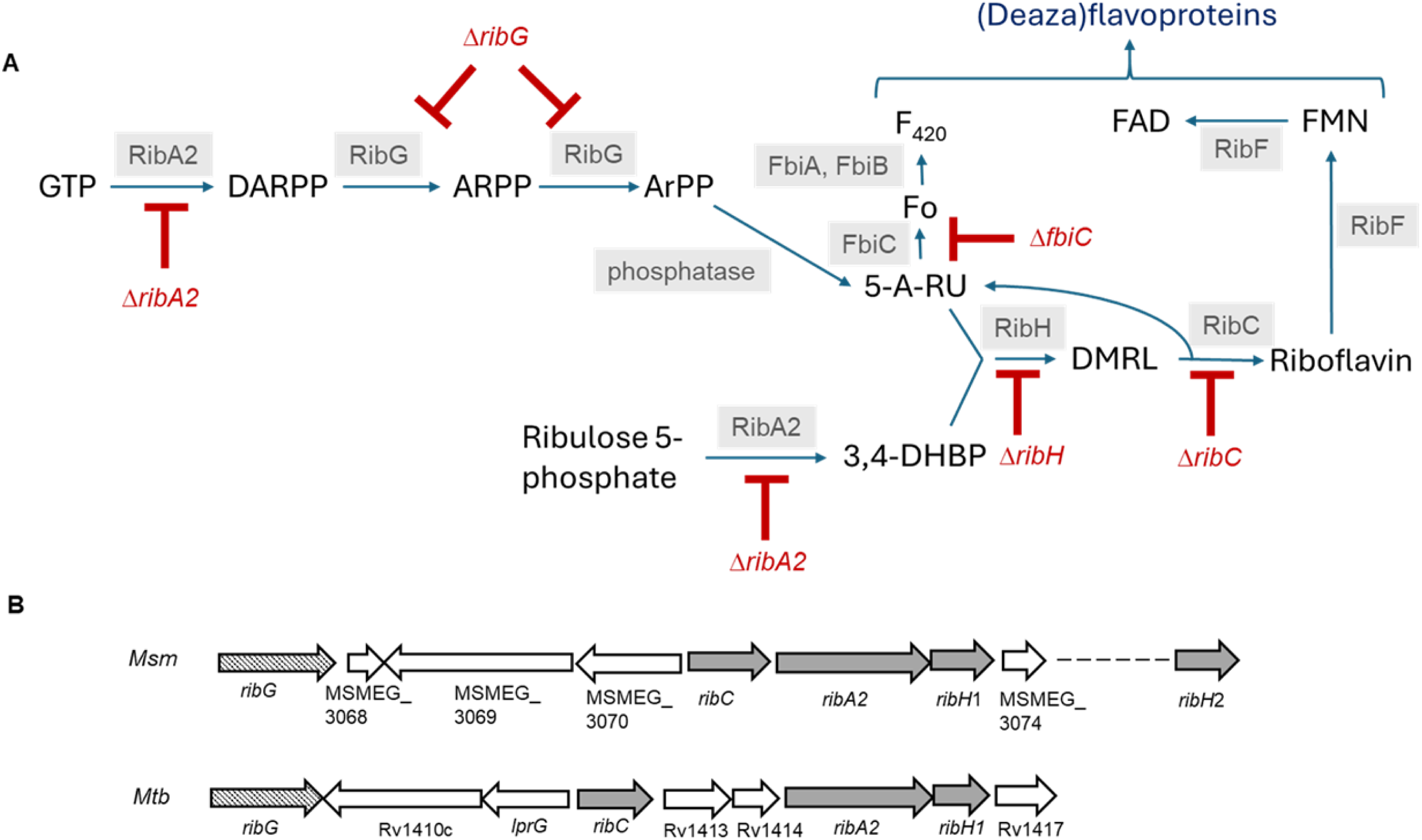
Riboflavin biosynthesis and utilization pathways in mycobacteria. **(A)** In the biosynthesis pathway, ribulose-5-P and GTP are converted to 3,4-dihydroxyl-2-butanone 4-phosphate (3,4-DHBP) and 2,5-diamino-6-(5-phospho-D-ribosylamino)-pyrimidin-4(3H)-one (DARPP), respectively, by the bifunctional GTP cyclohydrolase II/ DHBP synthase, RibA2. DARPP is deaminated and the side chain subsequently reduced by the bifunctional riboflavin deaminase/ 5-amino-6-(5-phosphoribosylamino) uracil reductase, RibG, to form 5-amino-6-ribitylamino-2,4(1H,3H)-pyrimidinedione-5-phosphate (ArPP). Dephosphorylation of ArPP by an unknown phosphatase generates 5-amino-6-D-ribitylaminouracil (5-A-RU). 5-A-RU together with 3,4-DHBP are condensed by the lumazine synthase, RibH (RibH1/RibH2), to yield 6,7-dimethyl-8-ribityllumazine (DMRL). Two molecules of DMRL are converted to riboflavin and 5-A-RU by the riboflavin synthase, RibC via a dismutation reaction. In the utilization pathway, riboflavin is converted to FMN and FAD by the bifunctional kinase/ FAD synthetase, RibF. The deazaflavin, F_420_, is produced from 5-A-RU by the sequential action of FbiC, FbiA and FbiB. The deletion mutations are shown in red. (**B)** Genomic context of the riboflavin biosynthesis genes in Msm and Mtb.

A notable difference between Msm and Mtb is the redundancy in lumazine synthase activity in Msm where this function is served by a canonical RibH (MSMEG_3073, designated herein as RibH1) – the homolog of the sole RibH in Mtb (Rv1416) – and a second homologue, MSMEG_6598, designated here as RibH2 [17]. In prior work, we used a panel of inducible CRISPRi hypomorphs in genes involved in riboflavin biosynthesis and utilization to confirm the essentiality of the riboflavin pathway for mycobacterial growth and survival *in vitro* and demonstrate that Mtb and Msm can transport and assimilate exogenous riboflavin [17]. However, residual expression of target genes in induced hypomorphs could confound the ability to differentiate the contributions of individual riboflavin pathway intermediates in MAIT cell recognition of mycobacteria. To avoid this potential complication, we created a Msm and Mtb riboflavin auxotrophs carrying deletions of individual genes in the riboflavin biosynthesis pathway for use in this study (**S1 Fig**). Importantly, the deletion mutations in *ribA2* eliminated both GTP cyclohydrolase II and DHBP synthase domains of RibA2, which catalyze the first and penultimate steps of the pathway, respectively.

The growth phenotypes of the Msm and Mtb mutants and their complemented counterparts were assessed in liquid and on solid media, with or without riboflavin supplement **(Fig 2 and S2 Fig)**. The Msm Δ*ribA2*, Δ*ribG*, Δ*ribH2*Δ*ribH1* and Δ*ribC* mutants, and the Mtb Δ*ribA2*, Δ*ribH* and Δ*ribC* mutants were auxotrophic for riboflavin whereas the Msm Δ*fbiC* mutant showed no growth phenotype. Limited complementation of the growth kinetics in liquid culture was observed for the Msm Δ*ribA2*, Δ*ribH1* and Δ*ribH2*Δ*ribH1* strains (complemented with *ribH2* or both) whereas restoration of growth approximating the corresponding wildtype was observed for the other complemented strains. The riboflavin auxotrophy of the Mtb Δ*ribH* and Msm Δ*ribH2*Δ*ribH1* mutants confirmed that under the conditions tested, these organisms depend on the enzymatic production of DMRL by lumazine synthase and as such, this cellular function cannot be served by non-enzymatic production of this pathway intermediate [21].

**Fig 2.**
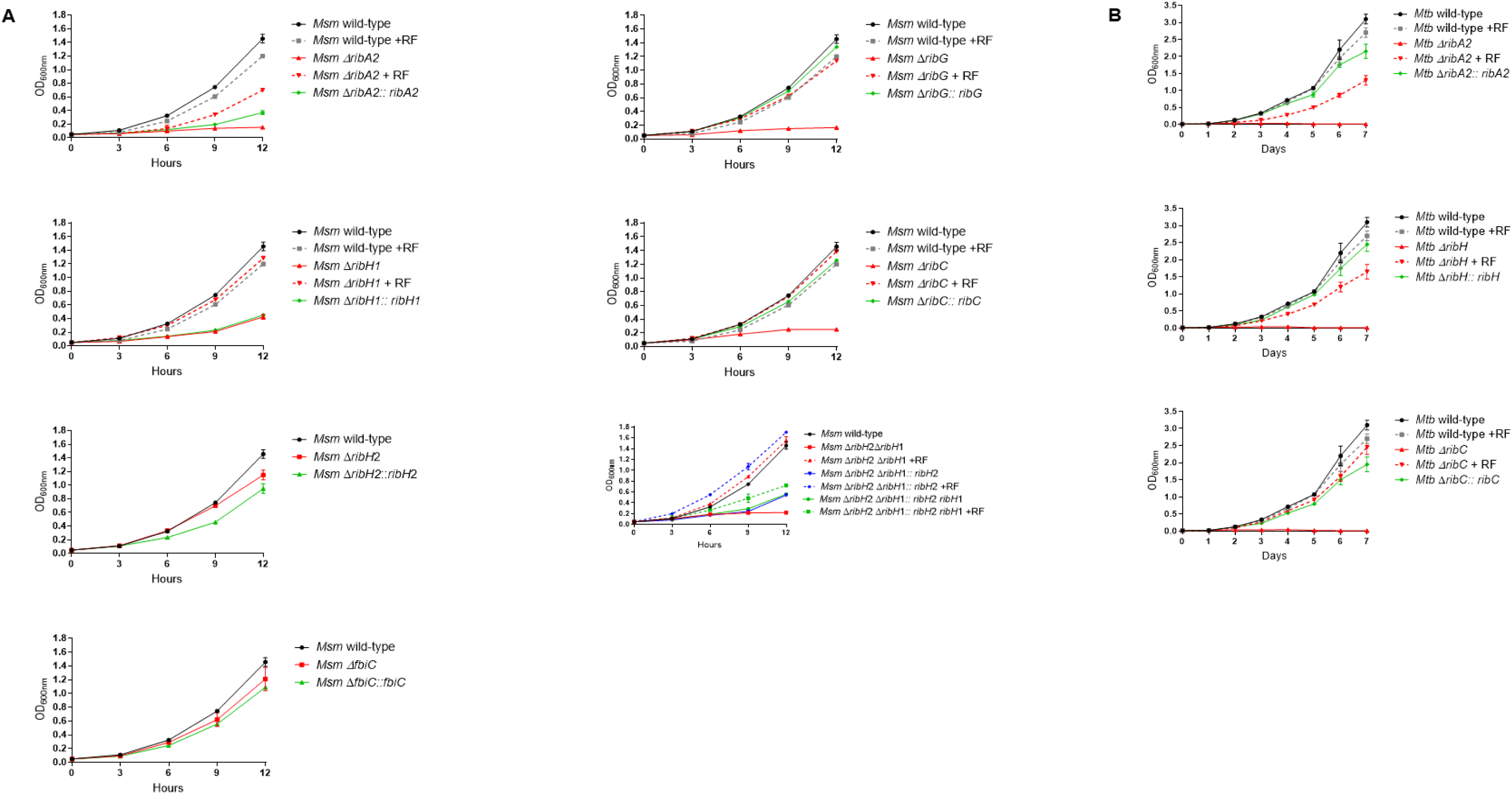
Growth kinetics of wildtype, knockout and complemented mutant strains of Msm (A) and Mtb (B) in 7H9 media. Error bars represent standard deviation from two biological replicates. Statistical comparisons, against wild-type were performed using a one-way ANOVA and Dunnett’s multiple comparison test whereby statistical significance is represented by *p* < 0.005, shown by an asterisk.

Riboflavin pathway transcript and protein levels in the Msm wildtype, knockout and complemented strains were measured by qRT-PCR and Data Independent Acquisition (DIA) proteomics, respectively (**Fig 3**). As the auxotrophs required riboflavin for growth, we assessed the impact of riboflavin supplement on the levels of riboflavin pathway transcripts and proteins in wildtype Msm and found no significant effects. The absence of the cognate transcript and protein was confirmed in the knockout mutants with variable restoration thereof in the complemented derivatives in a manner consistent with the corresponding growth phenotypes (**Fig 2 and S2**). Certain knockouts also affected the expression of other members of the pathway, e.g., a 3-fold increase in *ribH2* transcript and RibH2 protein was observed in the Msm Δ*ribH1* mutant suggesting a regulatory interdependence of expression between the two lumazine synthases **(Fig 3G and H).** However, the Δ*ribH1* mutant was still growth impaired relative to wildtype **(Fig 2)** indicating that RibH2 cannot fully compensate for the loss of RibH1 in Msm, which suggests that RibH1 is the main catalytic enzyme for the production of riboflavin. We observed some differences in the effect of pathway gene knockout *vs*. knockdown on expression of other genes in the pathway, *e.g*., silencing of *ribH1* led to an upregulation of *ribG* and *ribF*, and silencing of *ribA2* suppressed the expression of *ribG*, *ribC* and *ribH1* in Msm [17] but such effects were not observed by knockout of *ribH1* or *ribA2.* The differences may be due in part to the fact that the target proteins were only partially depleted in the knockdowns under the conditions of the expression analysis [17] but are entirely absent in the knockouts.

**Fig 3.**
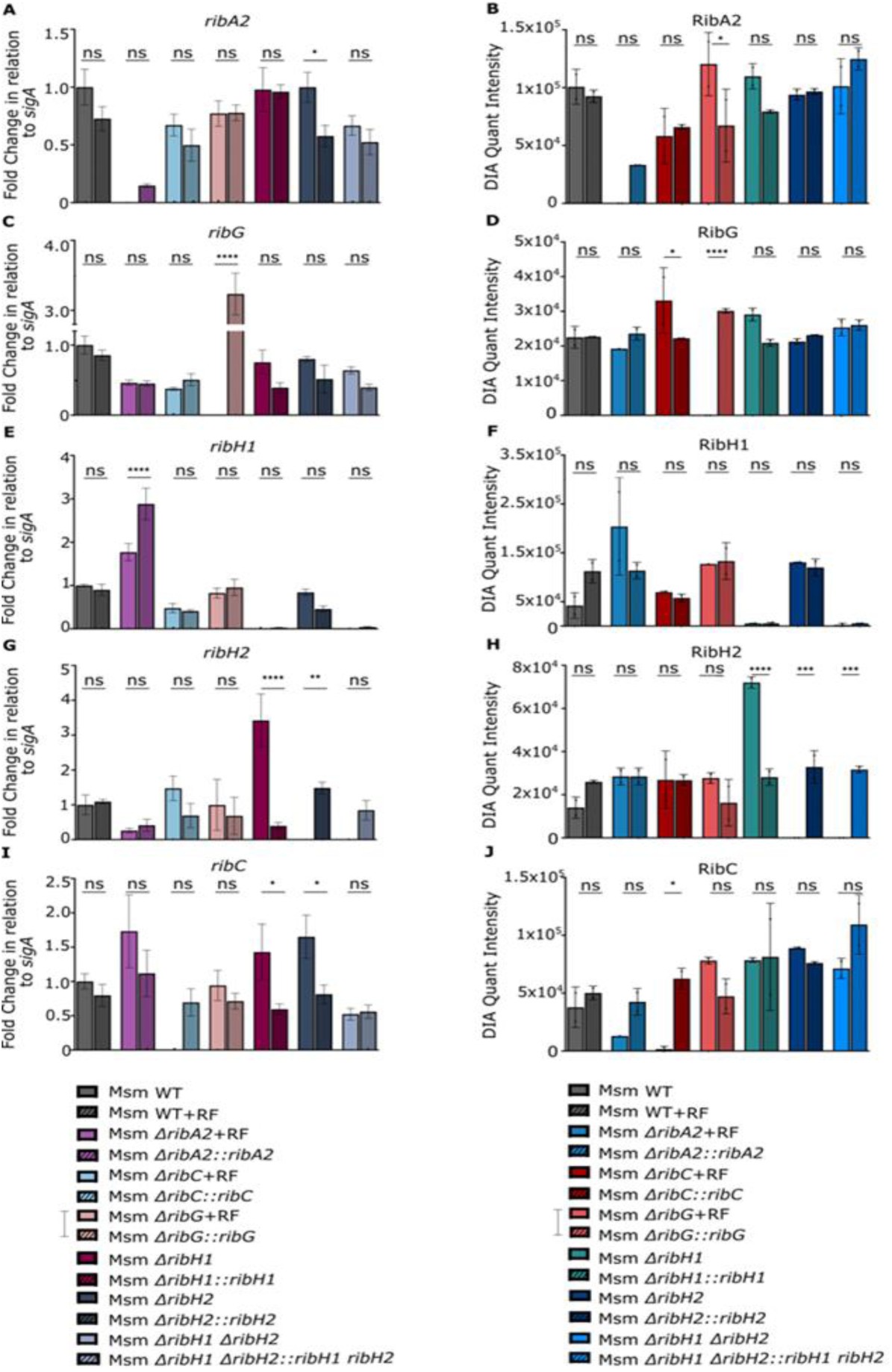
Quantification of riboflavin pathway transcript protein levels in Msm strains. **(A, C, E, G and I)** Fold change in *sigA-*normalized transcript levels relative to wildtype. **(B, D, F, H and J)** Protein abundance was measured using DIA. Data are shown as mean quantification ± SEM from two biological replicates. Statistical comparisons were performed using a one-way ANOVA and Sidak’s multiple comparison test whereby statistical significance is represented by p<0.05, p<0.001, p<0.0005 p<0.0001, shown by *, **, ***, **** respectively.

The Mtb Δ*ribA2*, Δ*ribH* and Δ*ribC* mutants likewise showed riboflavin auxotrophy with riboflavin-independent growth restored by genetic complementation. Knockout and complementation of *ribA2, ribH* and *ribC* were further validated by transcript and protein quantification (**Fig 4**). Additionally, stable isotope-labeled standards were made for peptides derived from RibA2, RibC and RibH with the goal of determining absolute peptide concentration and, by inference, the concentration of the intact protein using parallel reaction monitoring (PRM) (**S3 Fig**). Riboflavin supplement suppressed expression of *ribC* and enhanced expression of *ribA2* in wildtype Mtb **(Fig 4A and E)** but had no significant impact on expression of other pathway genes. Knockout of other pathway genes had no effect on RibA2 abundance but there was a reduction in expression of *ribA2* in the Δ*ribC* mutant which was restored by complementation **(Fig 4A and B)**. RibH transcript and protein levels were markedly lower than wildtype in the complemented Mtb Δ*ribH* strain **(Fig 4C and D)** but still sufficient to restore growth to wildtype levels **(Fig 2)**. In contrast, *ribH* transcript and RibH protein levels were increased 12-to-16-fold in the Δ*ribA2* mutant and its complement vs. wildtype **(Fig 4C and D)**, a finding confirmed by absolute quantification of RibH using PRM and likely due to the presence of a vector-borne promoter/s upstream of *ribH* in the Δ*ribA2* mutant (**S1 Fig**).

**Fig 4.**
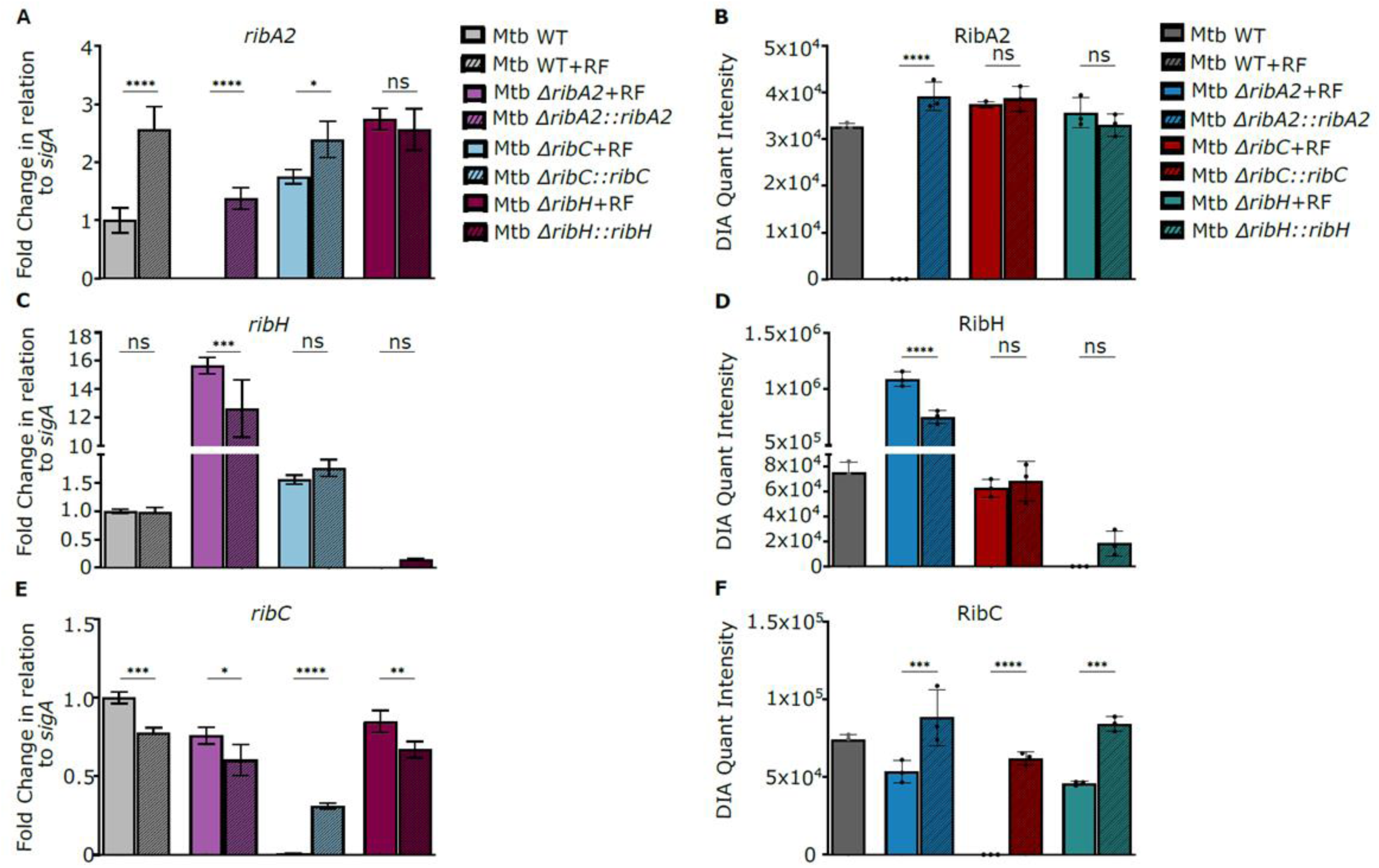
Quantification of riboflavin pathway transcript protein levels in Mtb strains. **(A, C, and E)** Fold change in *sigA-*normalized transcript levels relative to wildtype. Data are shown as mean quantification ± SEM from three biological replicates. **(B, D, and F)** Protein abundance was measured using DIA. Statistical comparisons were performed using a one-way ANOVA and Sidak’s multiple comparison test whereby statistical significance is represented by p<0.05, p<0.001, p<0.0005, p<0.0001, shown by *, **, ***, **** respectively. ns, not significant.

### Deletion of *ribA2* or *ribG* selectively ablates MAIT cell activation by Msm

To assess the impact of disrupting the riboflavin biosynthesis pathway on MAIT cell activation, Msm cultures grown in the presence of riboflavin were used to assess the ability to activate a panel of four MR1 restricted T cell clones: D426-G11, D481-C7 and D481-F12, TRAV1-2+ MAIT cell clones with varied TCRα chains **(S1 Table)** [8], and D520-E10, a TRAV1-2-MR1T clone [8]. These were selected to reflect the diversity of ligands seen by MR1Ts with D481 C7 uniquely able to detect the photolumazine antigens derived from Msm [8]. Each clone was tested by IFN-γ ELISPOT for its ability to recognize dendritic cells infected with mycobacteria at an MOI of 10 (**Fig 5**). Deletion of *ribA2* or *ribG* resulted in an 82-99% reduction in T cell recognition of Msm for all clones tested – a phenotype reversed by complementation. In contrast, deletion of *ribC* had no significant impact on T cell activation by Msm (**Fig 5**). Deletion of *ribH1* and *ribH2* alone or in combination had no impact on recognition of Msm by the three TRAV1-2+ clones, D426-G11, D481-C7 and D481-F12, whereas recognition of the TRAV1-2-clone, D520-E10, was reduced compared to wildtype response. However, this effect was not reversed by complementation, presumably due to the lack of complementation of *ribH1* deficiency revealed by growth phenotyping and expression analyses of the Δ*ribH1* and Δ*ribH2*Δ*ribH1* mutants (**Fig 2 and Fig 3**). Nonetheless, the diminished response of this clone, which uniquely recognizes Msm, implies that the antigen recognized by this clone could be a lumazine.

**Fig 5.**
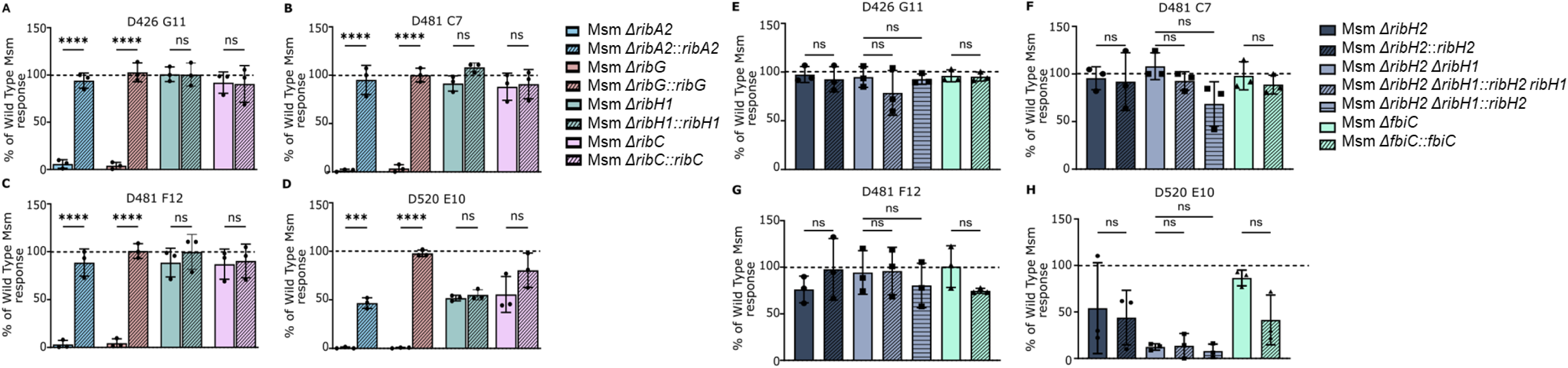
Impact of riboflavin pathway mutations on MR1T recognition of Msm. MR1T cell clone (1e4) IFN-γ response to DCs (1e4) incubated with Msm at a MOI of 10. Response was normalized to wildtype Msm. Data are representative of n=3 independent experiments. Statistical comparisons were performed using a one-way ANOVA and Sidak’s multiple comparison test whereby statistical significance is represented by p<0.05, p<0.001, p<0.0005 p<0.0001, shown by *, **, ***, **** respectively.

In addition to its role as a substrate of RibH and precursor of DMRL, 5-AR-U is also required for the production of the deazaflavin MAIT cell antagonist, F_0_ [18], via the action of FbiC. We therefore assessed the effect of blocking this route of 5-A-RU metabolism on MAIT cell activation using a *fbiC* deletion mutant of Msm. The Δ*fbiC* mutation had no effect on Msm viability (**Fig 2**) or recognition by the TRAV1-2+ T cell clones, D426-G11 and D481-C7 (**Fig 5**). However, there was a trend towards reduced recognition of Msm by the TRAV1-2+ clone, D481-F12, and the TRAV19-1 clone, D510-E10 (**Fig 5**). This again might suggest preferential recognition of the lumazine antigens found in Msm.

### Opposite effects of *ribA2* vs. *ribH* deletion on MAIT cell activation by Mtb

The Mtb strains were tested for their ability to activate three of the MR1 restricted T cell clones described above, namely, D426-G11, D481-C7, and D481-F12 (**Fig 6**). The TRAV1-2-clone D520-E10 was not used in this experiment as it is unable to recognize Mtb **(S4 Fig)**. Each clone was tested by IFN-γ ELISPOT for its ability to recognize the BEAS-2B bronchial epithelial cell line infected with Mtb at an MOI of 10. As the strains varied in their ability to activate the MAIT cell clones, we normalized these responses to the HLA-B45, Mtb8.4-specific clone [22]. As observed in Msm, the deletion of *ribA2* in Mtb reduced T cell recognition of all clones by 87-97%, an effect restored by complementation. On the other hand, the deletion of *ribH* significantly enhanced recognition of Mtb by all three MAIT cell clones (**Fig 6**), a different trend compared to the response observed in the single *ΔribH1*, *ΔribH2* or double *ΔribH2 ΔribH1* Msm mutants. Furthermore, knockout of the gene *ribC* had no significant effect on T cell recognition of Mtb.

**Fig 6.**
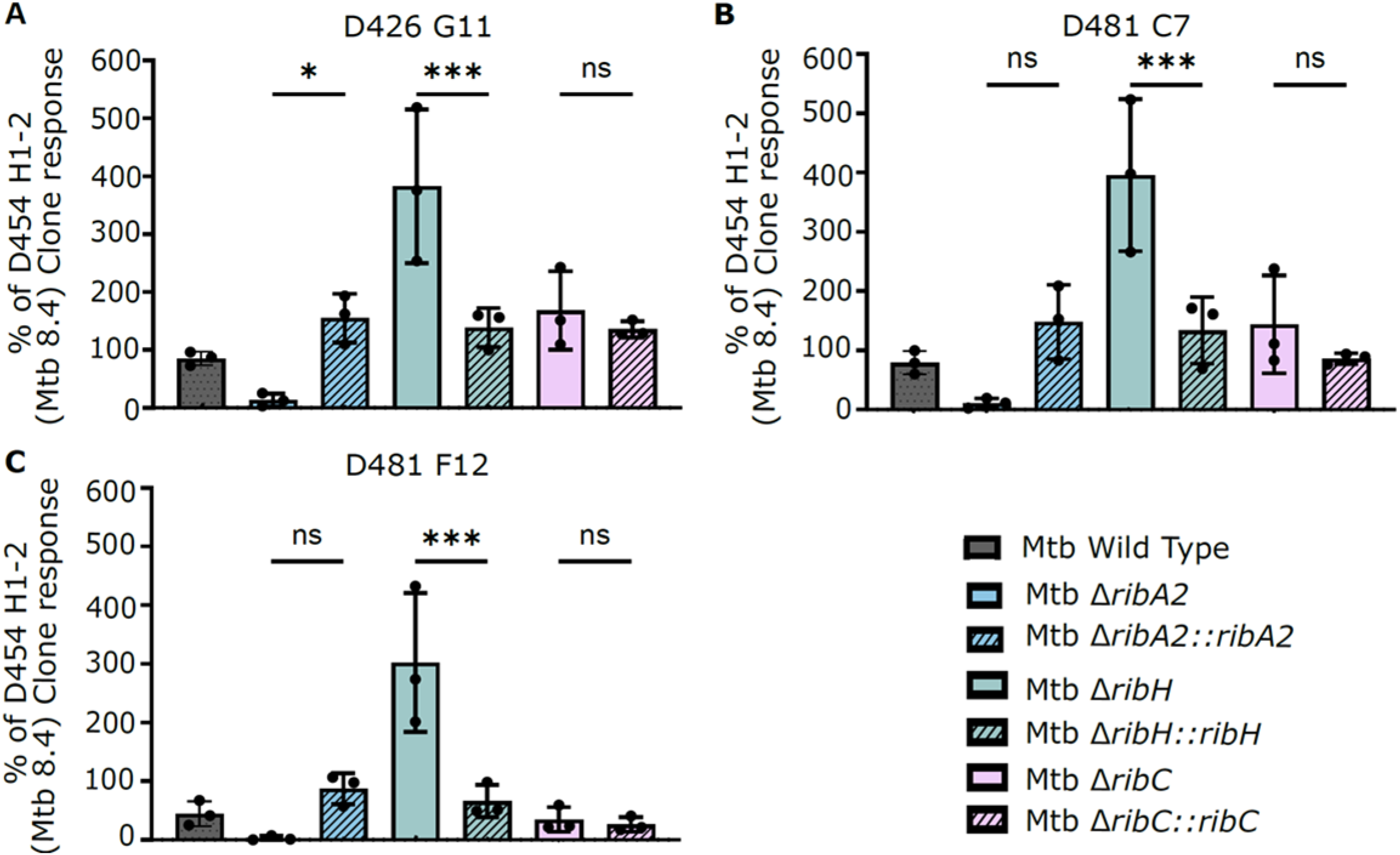
Differential impacts of riboflavin pathway knockouts on MR1T recognition of Mtb. The data represent the MR1T cell clone (1e4) IFN-γ response to DCs (1e4) infected at an MOI of 10 of the indicated Mtb strain. The response for each Mtb strain was normalized to the response of D454 H1-2, a classically restricted CD8 T cell clone that recognizes Mtb8.4 [22]. Data are representative of n=3 independent experiments. Statistical comparisons were performed using a one-way ANOVA and Sidak’s multiple comparison test whereby statistical significance is represented by p<0.05, p<0.001, p<0.0005 p<0.0001, shown by *, **, ***, **** respectively.

## Discussion

In this study, we investigated the association between riboflavin biosynthesis and MAIT cell activation by mycobacteria using strains of Msm or Mtb in which riboflavin biosynthesis was blocked at the early (*ribA2* or *ribG*), penultimate (*ribA2* or *ribH*) or last step of the pathway (*ribC*) (**Fig 1**). Deletion of *ribA2* or *ribG* blocks the production of 5-A-RU and the downstream metabolites, DMRL and riboflavin, whereas loss of RibH function by deletion of *ribH* in Mtb or *ribH1* and *ribH2* in Msm only blocks the production of DMRL and riboflavin, whereas *ribC* deletion only blocks riboflavin production.

The riboflavin pathway produces both MAIT cell agonists and antagonists which differ in terms of MR1 binding affinity and potency [23]. These include intermediates (F_0_, DMRL) and the end-product (riboflavin) of the pathway produced by enzyme-catalyzed reactions and secondary metabolites formed by non-enzymatic reactions of pathway intermediates with molecules from other pathways, which, in the case of Mtb, could include those derived from the host (**Fig 7**). The level of activation of a given MR1T is therefore determined by its selectivity for MR1 ligand recognition and balance between the level of agonists vs. antagonists. While we have found that both F_0_ and riboflavin are relatively weak antagonists [8], it is possible that the metabolic state of the microbe could influence the relative proportion of these ligands. The latter depends, in turn, on the metabolic state of the organism under the conditions tested. For example, the level of the toxic aldehyde, methylglyoxal, produced endogenously by Mtb [24] and also by host macrophages [25], from metabolites such as dihydroxyacetone phosphate could influence the production of 5-A-RU derived ligands. Likewise, an increased demand for F_420_ for persistence of Mtb under the conditions encountered by Mtb during infection (hypoxia, oxidative stress, nitrosative stress) and/or under drug pressure [18, 26] could drive the metabolic flux of 5-A-RU in mycobacteria towards the production of deazaflavins at the expense of lumazine agonists. Therefore, the agonist/ antagonist balance, and the derangement thereof by perturbing the metabolic flux through the biosynthetic pathway, could only be inferred indirectly by comparing the phenotypic readout of MR1T activation by wildtype vs. mutant strains grown under standard *in vitro* culture, albeit supplemented with riboflavin.

**Fig 7.**
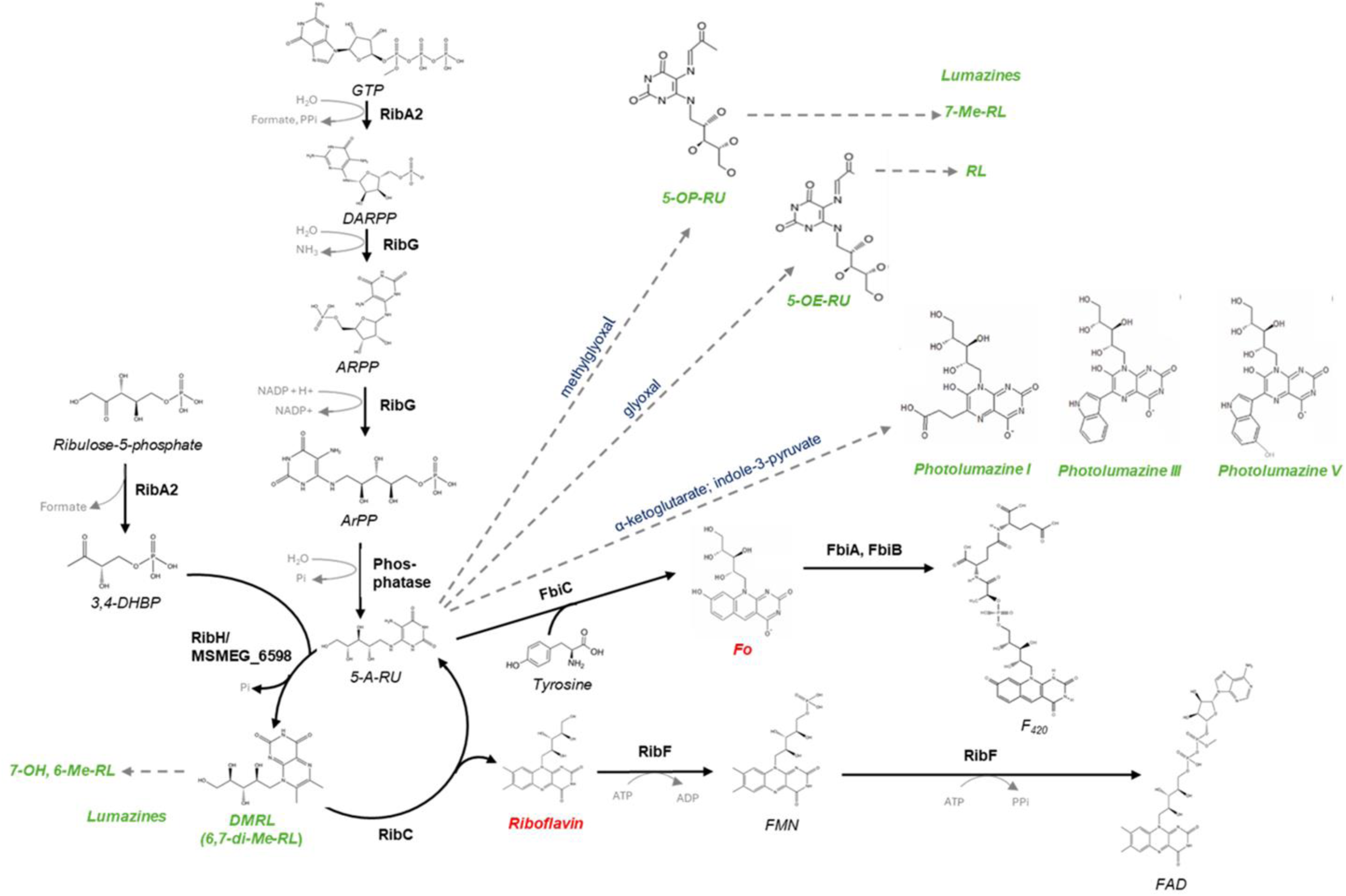
MAIT cell agonists and antagonists produced via the riboflavin pathway in mycobacteria. Solid black arrows denote enzyme-catalysed reactions. Dashed grey arrows denote non-enzymatic reactions. MAIT cell agonists (green) or antagonists (red) are shown in bold italics. RL, ribityllumazine. The production of photolumazines has only been observed in Msm and not in Mtb [8] [9].

We observed both similarities and differences between the two mycobacterial species in terms of the impact of riboflavin pathway disruption on MAIT cell activation. On the one hand and as observed in other microbes [7] – blocking the formation of 5-A-RU by disrupting the pathway at an early stage had a major impact on MAIT cell activation, confirming the centrality of this pathway intermediate for production of mycobacterial MR1T agonists. Moreover, the profoundly reduced ability to recognize Msm incapable of producing 5-A-RU applied equally to TRAV1-2+ and TRAV1-2-MR1Ts. However, residual MAIT cell activation was observed for Msm Δ*ribA2*, Msm Δ*ribG* and Mtb Δ*ribA2* thus implicating an alternate source/s of minor mycobacterial MAIT cell agonists distinct from the riboflavin pathway. Furthermore, for both Msm and Mtb, activation of all MR1Ts tested was unaffected by knockout of *ribC* which blocks the enzymatic route of DMRL metabolism to produce riboflavin and 5-A-RU. On the other hand, deletion of *ribH* enhanced MAIT cell recognition of Mtb by all three clones, D426-G11, D481-C7 and D481-F12 by 3-4-fold whereas in Msm, the Δ*ribH2*Δ*ribH1* mutant was recognized as well as the parental wildtype by these clones. Thus, blocking the enzymatic production of the MAIT cell agonist, DMRL, did not adversely affect MAIT cell recognition in either Mtb or Msm, and paradoxically, enhanced it in Mtb, possibly due to enhanced production of 5-A-RU-associated activating ligands. Conversely, this phenotype was not recapitulated in Msm (**Fig 7**) [9]. That Msm also preferentially produces lumazine antigens is suggested by the observation that the ablation of FbiC impacted the recognition of Msm by the non-TRAV1-2 T cell clone D520 E10, which recognizes Msm and not Mtb.

Other studies have demonstrated variable impacts of modulating flux through the riboflavin biosynthetic pathway on MAIT cell activation. Using an alternative approach involving overexpression rather than knockout of riboflavin pathway genes, Dey *et al*. also observed marked differences between mycobacterial species [16]. In that study, Mtb *ribH* overexpression modestly reduced activation of D426-G11, D481-C7 and D481-A9 by Mtb CDC1551 but had the opposite effect in Msm, enhancing activation of these MR1Ts up to ∼ 4-fold in an MOI-dependent manner. Mtb *ribH* overexpression also enhanced by 9.3-fold the activation of D426-G11 by *M. bovis* BCG, which, unlike Mtb and Msm (and for reasons that are unclear) could not tolerate overexpression of *ribA2*, *ribG* and *ribF* [16]. In *S*. *typhimurium* ST313 lineage 2 strains, evasion of MAIT cell recognition was shown to be attributable to overexpression of DHBP synthase, which produces the second substrate for RibH [15].

In summary, the data presented herein support a central role for the riboflavin pathway, and specifically 5-A-RU in generating a diverse array of mycobacterial ligands, which in turn are associated with selective T cell recognition. The antigen 5-OP-RU was initially described for both *E. coli*, and *Salmonella* species, and hence supported the concept that MAIT cells were innate in their use of a highly conserved presentation molecule (MR1), a conserved TCRα chain (TRAV1-2). However, in our previous work with both Msm and Mtb, we have observed a diversity of both MR1-bound ligands and activating fractions [8]. Specifically, in our ongoing work we have not observed the lumazine antigens seen in Msm in Mtb, and specifically note that we have not observed DMRL in both mycobacterial specificities. While the failure to observe 5-OP-RU or DMRL could reflect their chemical instability, it does not account for the diversity of ligands that we have observed. With regards to TCR usage, we have found that the selective recognition of mycobacterial antigens is associated with TCRβ chain usage [27]. Hence this work supports the hypothesis that further identification of these antigens could be used to define the extent to which these T cell responses are associated with infection with Mtb or could be harnessed for an improved vaccine. We also note that while ablation of RibA2 has a profound effect on the recognition of both Msm and Mtb, there were residual activating antigens in both these strains. Hence, it is likely that there remain non-5-A-RU dependent antigens yet to be discovered.

## Materials and methods

### Media, culture conditions, strains, plasmids and chemicals

The plasmids used and constructs generated in this study are described in **S2 Table**. All strains generated were derivatives of either *E. coli* DH5α, Msm mc^2^ 155 or Mtb H37RvMA, shown in **S3 Table**. *E. coli* DH5α was grown in Luria-Bertani (LB) medium (10g/L tryptone, 5g/L yeast extract and 10g/L NaCl) and plated onto Luria-Bertani agar (LB agar), containing LB media components supplemented with 15g/L agar. Mycobacterial strains were grown in Middlebrook 7H9 OADC medium (4.7 g/L 7H9 (Millipore Sigma), 2g/L glycerol (Millipore Sigma), 2.5 ml 25% Tween 80, and 100 ml of oleic acid-albumin-dextrose-catalase (OADC) enrichment (Millipore Sigma). Cells were plated onto Middlebrook 7H10 agar plates (19g/L 7H10 (Millipore Sigma), 100 ml OADC and 5 ml glycerol). Msm was grown in Erlenmeyer flasks in a rotating incubator set at 100 rpm at 37°C and unless otherwise stated Mtb was cultured in sealed cell culture flasks with no agitation at 37°C. Where required 7H9 broth or 7H10 solid media was supplemented with 50 µg/ml kanamycin, 50 µg/ml hygromycin or 2.5 µg/ml gentamicin. Riboflavin auxotrophs were supplemented with exogenous riboflavin at a concentration of 83 µM for Msm or 21 µM for Mtb. Media used to select for knockout mutants constructed by allelic exchange [28] or ORBIT [29] are described below.

### Construction of mutant and complemented strains

Allelic exchange by homologous recombination was used to knockout *ribC* in Mtb and *ribC*, *fbiC* and MSMEG_6598 in Msm, as described [28]. Briefly, 1000-2000 bp fragments amplified up- and downstream of the gene of interest were ligated and cloned in p2NIL. The *Pac*I cassette from either pGOAL19 or pGOAL17 cloned in the *Pac*I site in the p2NIL **(S2 Table).** The resulting suicide construct was electroporated into the mycobacterial strain and single crossover clones selected by plating onto 7H10 media supplemented with 50 µg/ml kanamycin, 50 µg/ml hygromycin and 40 µg/ml X-gal (for pGOAL19 constructs) or 50 µg/ml kanamycin and 40 µg/ml X-gal (for pGOAL17 constructs). Double crossover recombinants were selected by plating onto media supplemented with 2% sucrose, 40 µg/ml X-gal and riboflavin at concentration of 83 µM for Msm or 21 µM for Mtb. The ORBIT system [29] was used to create individual gene knockouts of *ribA2*, *ribG* or *ribH* in Msm and *ribA2* or *ribH* in Mtb. Briefly, a targeting oligonucleotide containing 70 bp of homology to the target gene flanked by the BxbI phage *attP* sequence **(S4 Table)** was co-electroporated with pKM464 into the mycobacterial strain carrying pKM461 and selected on 7H10 agar containing 10% sucrose and 50 µg/ml hygromycin. To create complementation vectors, ca. 300-400 bp upstream of the gene along with its open reading frame were amplified and cloned into an integrating vector. All primers and amplicons are described in **S4 Table**. The complementation vectors were introduced into the knockout mutants by electroporation and selected on the appropriate antibiotic selection plates. DNA was extracted from all gene knockout and complement strains using the CTAB method [30] and the gene deletion/ gene restoration verified by whole genome sequencing.

### RNA extraction

Mtb and Msm strains were inoculated in 80 ml of Middlebrook 7H9 OADC broth supplemented with 0.01% tyloxapol until mid-log phase (OD_600_ between 0.4 and 0.6) was reached. Media was supplemented with riboflavin as necessary. Afterwards, cells were pelleted, resuspended in 1 ml of TriZOL (Thermo Scientific) and 5 µl of glycogen (Thermo Scientific) and then transferred to screw-cap homogenization microtubes prefilled with Lysing Matrix B (MP Biomedical). The cells were homogenized using the FastPrep-24 bead beater (MP Biomedical) at 6 m/s for 30 seconds (6 rounds), cooling on ice for 2 min between cycles. After centrifugation at 12,000×*g* for 120s, homogenate was collected in a new tube and treated with 200 µl of chloroform followed by vortexing for 15 seconds and incubation at room temperature for 5 minutes. Samples were centrifuged and at 12,000×*g* for 15 minutes at 4°C and upper aqueous phase was transferred into new tubes containing 500 µl of cold isopropanol for overnight precipitation. Samples were centrifuged at 12,000×*g* for 10 min and supernatant disposed. Pellet was washed with 1 ml of cold 80% ethanol and then resuspended with nuclease free water. Extracted RNA was processed for DNase treatment and clean-up using the RNeasy Mini Kit (QIAGEN) and RNase free DNase Set (QIAGEN) according to manufacturer’s instruction to eliminate genomic DNA. RNA was quantified using a NanoDrop (Thermo Scientific) and Agilent TapeStation.

### Gene expression analysis

Two hundred ng of RNA was retrotranscribed to complementary DNA (cDNA) using the SuperScript IV VILO Master Mix (Thermo Scientific) according to manufacturer’s instruction. PowerTrack SYBR green Master Mix (Thermo Scientific) with 250 nM of primer pair was used to amplify the target region of interest for 5 ng of cDNA following manufacturer’s instruction. Quantification was run on the QuantStudio 3 Real-Time PCR System (Thermo Scientific). A no reverse transcriptase (no-RT) control was included for all samples to ensure that there was no genomic DNA contamination. Amplicon size was confirmed by melting curves for each sample. All qRT-PCR experiments were performed with two technical replicates derived from three biological replicates for Mtb and two biological replicates for Msm. Transcript levels of the target genes were normalized to the housekeeping gene *sigA* and calibrated to wild type Mtb and Msm to calculate the ΔΔCt and determine fold difference in gene expression. The primers used are listed in **S5 Table**.

### Growth kinetics of mycobacteria

Msm and Mtb growth kinetics in 7H9 media were monitored by OD_600_ as previously described [17].

### Growth of Mtb and Msm for untargeted and targeted proteomics

Glycerol stocks of Mtb and Msm strains were used to start cultures on 7H11 agar plates (Millipore Sigma, Cat No.: M0428) supplemented with the appropriate antibiotics with or without riboflavin. Plates were incubated for 2 weeks for Mtb and 4 to 5 days for Msm at 37°C. Cells were scraped from plates into 7H9 media supplemented with OADC with or without riboflavin and grown at 37°C until mid-log phase. Mtb cells from all strains were inactivated by γ-irradiation and inactivation was confirmed using the Alamar Blue assay. Cells were harvested and washed 3 times either with phosphate buffered saline (PBS) or water to remove media and stored at -80°C until further processing. Cell pellets were further processed as previously described [17].

### Liquid chromatography - MS/MS analysis

Thirty µg of retentate from whole cell lysate from Msm and Mtb after ultrafiltration with 3 kDa filters (Millipore Sigma, Product Number: UFC9003), was subjected to in-gel trypsin digestion as described previously [31]. Digested samples were resuspended in 120 µl of loading buffer (3% acetonitrile, 0.1% formic acid in water). All samples were analyzed using nano-UHPLC (ultrahigh-pressure liquid chromatography) nanoElute (Bruker Daltonics, USA) coupled to a timsTOF Pro (Bruker Daltonics, USA) mass spectrometer. 0.25 µg of total peptides were loaded onto a trap cartridge using PepMap™ Neo Trap Cartridge (Thermo Scientific, Cat: 174500) (5 µm C18 300 µm X 5 mm) packed with spherical, fully porous, ultrapure silica and reverse phase UHPLC was performed in a PepSep Ten Column (Bruker Daltonics, Part No:1895802) (1.5 µm C18 75 µm X 25 cm). Mobile phase A was 0.1% formic acid in water; mobile phase B was 0.1% formic acid in acetonitrile. Liquid chromatography (LC) separation was carried out at a flow rate of 0.5 µL/min using a linear gradient of 5% solvent B to 30% solvent B over a duration 17.8 minutes, then 30% solvent B to 95% solvent B for 2.5 minutes. Thereafter, 95% solvent B was held for 2.4 minutes for equilibration.

Mass spectra and tandem mass spectra were obtained using the parallel accumulation serial fragmentation (PASEF) acquisition method in positive ion mode. Captive spray source was operated with the following settings: 1600 V capillary voltage, 3.0 l/min of dry gas at 200°C with the NanoBooster switched off.

A SWATH-MS spectral library was generated by analyzing samples using data-dependent acquisition (DDA-PASEF). DDA was set to obtain high resolution MS scans over the m/z range from 100 to 1700 and ion mobility range from 0.70 Vs/cm^2^ to 1.50 Vs/cm^2^. Ramp time was set at 100 msec and accumulation time was set at 100 msec. The collision energy was set to follow a linear function starting from 20 eV for 0.6 V. s/cm^2^ to 59 eV for 1.6 V. s/cm^2^. The number of PASEF MS/MS was set to 6. For DIA-PASEF analysis, data was acquired with a method consisting of 24 mass steps per cycle **(S6 Table)**. Mass width was set at 50 Da with 1 Da overlap between windows covering a mass range between 307.8 to 1484.8 Da. The ion mobility range (1/K_0_) covered was between 0.71 to 1.44. The collision energy was set to follow a linear function starting from 20 eV for 0.6 V. s/cm^2^ to 59 eV for 1.6 V. s/cm^2^.

### Untargeted proteomics analysis by Data Independent Acquisition (DIA)

DIA-PASEF data was processed on FragPipe software v. 19.1 (MSFragger version 3.7, IonQuant version 1.8.10, Philosopher version 4.8.1) using the DIA_SpecLib-Quant workflow as described [32]. DIA raw files were loaded with DIA Quant selected as data type while DDA raw files were loaded for spectral library generation. A search was conducted using default settings with the FASTA sequence obtained from the Mycobrowser database (Release 4, 2021-03-23). Decoy and contaminant sequences were generated by FragPipe. Default settings of MSFragger was used, supplemented with oxidation of M (+ 15.9949 Da mass shift) acetylation at N-terminal (+ 42.0106) as variable modifications. Advanced output options were set to write calibrated mzML and MGF files. Tryptic searches were performed with 20 ppm precursor and fragment tolerance permitting up to 2 missed cleavages. Search results were filtered to 1% FDR at both protein and peptide levels using peptide prophet. Output files with quantification based on summed intensity of all ions for a single peptide and summed intensity of peptides for a single protein were generated. The output file containing summed intensity of peptides for a single protein was used for quantification of proteins and is provided in **S6 Table.**

### Targeted proteomics analysis by parallel reaction monitoring - PASEF Mass Spectrometry (PRM-PASEF MS)

DDA peptide search results from MSFragger version 3.7 were used to build a spectral library in Skyline [33]. The Ion Mobility library was generated on Skyline software from raw DDA files. The protein list was trimmed to contain only proteins from the riboflavin biosynthetic pathway. Tryptic peptides were filtered to have length only be between 8 to 30 amino acids.

Tryptic peptides containing C, M, D-P, and W were excluded. Peptides for each protein were further screened based on the shape of the precursor chromatogram and absence of the precursor peak from the cognate knock-out sample. Ion types were limited to only precursor ions (2 and 3 ion charges) and 6 y ions (1 and 2 ion charges). The retention time window was set at 3 minutes and mass accuracy set at 10 ppm for both MS1 and MS2 filtering. The method was exported to timsTOF engine (Bruker Daltonics, USA) as an unscheduled method. 0.25 µg of each Mtb sample was loaded for reverse phase UHPLC and MS. performed. Analytical settings were kept the same as previously described (LC-MS/MS Section) with the inclusion of a target list containing the Mtb peptides to quantify in the PRM-PASEF method generated.

Raw data was exported into Skyline software and peptides with the best peak shape and intensity were selected for further optimization. At least, one peptide per protein was selected for the final method. For absolute quantification, peptides labeled with heavy C-terminal Lysine (K) or Arginine (R) were purchased (Biosynth Ltd, UK) to be used as internal standards. Limit of detection (LOD) and Limit of quantification (LOQ) were calculated using a linear regression method as described elsewhere [34]. Details of the peptide sequences and transition list used for the PRM-PASEF experiment are shown in **Table S7**.

### Absolute quantification of riboflavin pathway proteins by PRM-PASEF MS

A mix of the heavy labeled peptides containing 10^-1^ – 10^-5^ pmol/µl of each peptide was used to resuspend each of the digested samples at a final concentration of 250 ng/µl of digested protein **(S8 Table)**. After resuspension, the sample peptide concentration was estimated using a Nanodrop 1000c and the concentration was adjusted to ensure equal loading of all samples.

The developed PRM-PASEF MS method was exported as a scheduled method and used to obtain the Ratio to Heavy Standard for each of the monitored peptides. Briefly, raw data from each injection was imported into Skyline [33] and peak boundaries for each peptide were manually validated and adjusted if necessary. The Ratio to Heavy Standard for each peptide as calculated by Skyline was exported into Prism GraphPad for statistical analysis.

### Whole genome sequencing

Genomic DNA was analyzed at the Genomics and Microarray Core at the University of Colorado Anschutz using a NovaSEQ 6000 Instrument, paired end 150 cycles 2x150. Reads were subjected to quality control using FastQC before and after trimming. Trimming was performed using trimmomatic. Alignment of reads to the Mtb H37Rv genome (NC_000962.3) or Msm mc^2^ 155 (CP000480.1)/ (NZ_CP054795.1) was performed using Bowtie2. Due to the high density of reads, the aligned sam files were subjected to subsample using samtools. Samtools were also used to determine coverage results. Subsample alignment was used to determine SNPs using lofreq with a minimum coverage of 10 and minimum frequency of 80. The bam file was visualized using the Geneious software. The expected deleted regions were confirmed by visual inspection and very few inadvertent second-site mutations were detected in the deletion mutants of Msm and Mtb relative to their parental wildtype strains (**Table S9A, B1, B2 and B3**).

### T cell clones

All T cell clones were expanded and maintained as previously described [4, 22]. Five T cell clones were used in this work. All have been published previously: D426 G11, D481 C7, D481 F12 - MAIT cell clones expressing TRAV1-2 [4, 8]; D520 E10 is a MR1-restricted T cell clone expressing TRAV19-1[8], and D454 H1-2 is a CD8 T cell clone that recognizes Mtb8.433-43 and is restricted by HLA-B*15:01 [22] (see **S9 Table** for details of TCRs).

### IFN-γ ELISpot assays

All Msm and Mtb strains were grown in the presence of riboflavin with or without supplements as needed (see above) to an OD_600_ ∼ 0.5 and frozen down. For all ELISPOT assays, ELISPOT plates were coated with anti-IFN-γ antibody (Mabtech, clone 1-D1K, Cat No. 3420-3-1000, used at 10 μg/ml) as previously described [35]. After overnight incubation at 4°C, ELISPOT plates were washed three times with phosphate-buffered saline and then blocked with RPMI 1640 + 10% human serum for at least 1 hour. For experiments with Msm approximately 1x10^4^ DC were plated in the ELISPOT wells in RPMI+10% HuS. Msm bacilli were added to the wells at concentrations indicated for each experiment. PHA was used as positive control. After at least 1 hour, 1-2x10^4^ T cell clones were added, and the plate was incubated overnight at 37°C. IFN-γ ELISPOTs were enumerated following development as previously described [35]. For experiments with live Mtb, 5 × 10^5^ DC were plated in each well of an ultralow adherence 24 well tissue culture plate. Mtb was added at the MOI indicated and incubated overnight. 1x10^4^ DC were plated in the ELISPOT wells in RPMI+10% HuS. 1-2x10^4^ T cell clones were added, and the plate was incubated overnight at 37°C. IFN-γ ELISPOTs were enumerated following development.

### Statistical analyses

Statistical analysis was conducted by one-way ANOVA and either Sidak’s or Dunnett’s multiple comparison test. Statistically significant difference applied is stipulated in the figure legend. Statistical calculations were performed using GraphPad Prism 10 statistical software (GraphPad Software, Inc., San Diego, CA). For proteomics, qRT-PCR and the EliSPOT assay, statistical comparisons were performed using a one-way ANOVA followed by Dunnett’s or Sidak’s multiple comparison test. These analyses were performed using GraphPad Prism 9 and statistical significance was established at *p* < 0.05.

## Acknowledgements

This research was supported by an Oppenheimer Fellowship from the Oppenheimer Memorial Trust (to V.M.), and grants from the Bill and Melinda Gates Foundation (INV-004757), the Broad Institute (to V.M.), the South African Medical Research Council (to V.M.), the Department of Science and Innovation and National Research Foundation of South Africa (to V.M.), NIAID (R01AI147954, to D.M.L.), a subaward from the University of Chicago ((NIH NIAID R01AI147954, E.J.A., W.H. and D.M.L. MPIs)) (AWD100279, to K.M.D. and C.M.) and internal funds from Colorado State University (to K.M.D.).

## Author contributions

Conceptualization: Melissa Chengalroyen, Karen Dobos, David Lewinsohn, Deborah Lewinsohn, Valerie Mizrahi, Digby Warner

Data curation: Nurudeen Oketade, Melissa Chengalroyen, Gwendolyn Swarbrick

Formal analysis: Nurudeen Oketade, Carolina Mehaffy, Gwendolyn Swarbrick, Melissa Chengalroyen

Funding acquisition: Valerie Mizrahi, David Lewinsohn, Karen Dobos, Deborah Lewinsohn, Erin Adams, William Hildebrand

Investigation: Melissa Chengalroyen, Nurudeen Oketade, Gwendolyn Swarbrick, Carolina Mehaffy, Aneta Worley, Mabule Raphela, Megan Lucas

Methodology: Nurudeen Oketade, Gwendolyn Swarbrick, Luisa Nieto Ramirez, Carolina Mehaffy, Melissa Chengalroyen, Karen Dobos, David Lewinsohn, Valerie Mizrahi

Resources: Valerie Mizrahi, Karen Dobos, David Lewinsohn

Supervision: Luisa Nieto Ramirez, Valerie Mizrahi, Karen Dobos, Deborah Lewinsohn, David Lewinsohn

Visualization: Nurudeen Oketade, Gwendolyn Swarbrick, Melissa Chengalroyen, Carolina Mehaffy

Writing – original draft: Melissa Chengalroyen, David Lewinsohn, Nurudeen Oketade, Valerie Mizrahi

Writing – review & editing: Melissa Chengalroyen, Nurudeen Oketade, Luisa Nieto Ramirez, Megan Lucas, Karen Dobos, David Lewinsohn, Valerie Mizrahi

## Supporting information

**S1 Fig. Schematic representation of the genomic loci of the riboflavin biosynthesis pathway mutants of Msm (left) and Mtb (right).** Deletion mutants constructed by ORBIT carry a vector sequence which include a *hyg* resistance marker whereas the allelic exchange mutants are unmarked.

**S2 Fig**. **Growth phenotyping on solid media.** Growth of Msm (Left) and Mtb (right) riboflavin pathway mutants and complemented counterparts assessed by streaking on Middlebrook 7H10 with or without riboflavin (RF) supplement. Riboflavin supplement was used at a concentration of 83 μM for culturing Msm, or 21 μM for Mtb.

**S3 Fig. Protein abundance of Mtb strains assessed by PRM.** Stable isotope-labeled standard (SIS) peptides were selected for selected riboflavin pathway genes for targeted quantification of proteins using LC-MS) Ratio to heavy standard represents protein abundance of RibC **(A)**, and RibH **(B and C)** in different strains of Mtb. Data are shown as mean quantification ± SEM of three biological replicates relative to Mtb wild type (WT). Statistical comparisons were performed using a one-way ANOVA and Sidak’s multiple comparison test whereby statistical significance is represented by p<0.05, p<0.001, p<0.0005 p<0.0001, shown by *, **, ***, **** respectively. ns, not significant.

**S4 Fig. Recognition of D520 E10 by Mtb.** The data represent the MR1T cell clone (1e4) IFN-γ response to DCs (1e4) infected at an MOI of 10 of with WT Mtb or in response to the positive control, PHA. Data are representative of n=3 independent experiments.

**S1 Table. TCR sequence of MR1T cell clones**

**S2 Table**. **Plasmids used in this study**

**S3 Table. Strains used in this study**

**S4 Table. Oligonucleotides used to create knockouts and complemented strains**

**S5 Table. Primers used for qRT-PCR analysis**

**S6 Table. Window scheme used for DIA-PASEF acquisition**

**S7 Table. Peptides used for targeted proteomics by PRM-MS**

**S8 Table. Peptides used for targeted proteomics by PRM-MS and concentration used in final assay**

**S9A Table. Confirmation of riboflavin pathway mutants by WGS**

**S9B Tables. SNPs detected in Msm knockout mutants compared to wild type**

**S9 B1 & B2 Tables. SNPs detected in Msm mutants and not in wild type**

**S9 B3 Table. SNPs detected in Mtb mutants and not in wild type**

## References

1. Huang S, Gilfillan S, Cella M, Miley MJ, Lantz O, Lybarger L, et al. Evidence for MR1 antigen presentation to mucosal-associated invariant T cells. J Biol Chem. 2005;280(22):21183–93.

2. Treiner E, Duban L, Moura IC, Hansen T, Gilfillan S, Lantz O. Mucosal-associated invariant T (MAIT) cells: an evolutionarily conserved T cell subset. Microbes Infect. 2005;7(3):552–9.

3. Sharma PK, Wong EB, Napier RJ, Bishai WR, Ndung’u T, Kasprowicz VO, et al. High expression of CD26 accurately identifies human bacteria-reactive MR1-restricted MAIT cells. Immunology. 2015;145(3):443–53.

4. Gold MC, Cerri S, Smyk-Pearson S, Cansler ME, Vogt TM, Delepine J, et al. Human Mucosal Associated Invariant T Cells Detect Bacterially Infected Cells. PLOS Biology. 2010;8(6):e1000407.

5. Le Bourhis L, Martin E, Peguillet I, Guihot A, Froux N, Core M, et al. Antimicrobial activity of mucosal-associated invariant T cells. Nat Immunol. 2010;11(8):701–8.

6. Kjer-Nielsen L, Patel O, Corbett AJ, Le Nours J, Meehan B, Liu L, et al. MR1 presents microbial vitamin B metabolites to MAIT cells. Nature. 2012;491(7426):717-23.

7. Corbett AJ, Eckle SB, Birkinshaw RW, Liu L, Patel O, Mahony J, et al. T-cell activation by transitory neo-antigens derived from distinct microbial pathways. Nature. 2014;509(7500):361-5.

8. Harriff MJ, McMurtrey C, Froyd CA, Jin H, Cansler M, Null M, et al. MR1 displays the microbial metabolome driving selective MR1-restricted T cell receptor usage. Sci Immunol. 2018;3(25).

9. Krawic JR, Ladd NA, Cansler M, McMurtrey C, Devereaux J, Worley A, et al. Multiple Isomers of Photolumazine V Bind MR1 and Differentially Activate MAIT Cells. J Immunol. 2024;212(6):933–40.

10. Meermeier EW, Laugel BF, Sewell AK, Corbett AJ, Rossjohn J, McCluskey J, et al. Human TRAV1-2-negative MR1-restricted T cells detect S. pyogenes and alternatives to MAIT riboflavin-based antigens. Nat Commun. 2016;7:12506.

11. Chen Z, Wang H, D’Souza C, Sun S, Kostenko L, Eckle SB, et al. Mucosal-associated invariant T-cell activation and accumulation after in vivo infection depends on microbial riboflavin synthesis and co-stimulatory signals. Mucosal Immunol. 2017;10(1):58–68.

12. Legoux F, Bellet D, Daviaud C, El Morr Y, Darbois A, Niort K, et al. Microbial metabolites control the thymic development of mucosal-associated invariant T cells. Science. 2019;366(6464):494-9.

13. Hartmann N, McMurtrey C, Sorensen ML, Huber ME, Kurapova R, Coleman FT, et al. Riboflavin Metabolism Variation among Clinical Isolates of Streptococcus pneumoniae Results in Differential Activation of Mucosal-associated Invariant T Cells. Am J Respir Cell Mol Biol. 2018;58(6):767–76.

14. Jahreis S, Bottcher S, Hartung S, Rachow T, Rummler S, Dietl AM, et al. Human MAIT cells are rapidly activated by Aspergillus spp. in an APC-dependent manner. Eur J Immunol. 2018;48(10):1698–706.

15. Preciado-Llanes L, Aulicino A, Canals R, Moynihan PJ, Zhu X, Jambo N, et al. Evasion of MAIT cell recognition by the African Salmonella Typhimurium ST313 pathovar that causes invasive disease. Proc Natl Acad Sci U S A. 2020;117(34):20717–28.

16. Dey RJ, Dey B, Harriff M, Canfield ET, Lewinsohn DM, Bishai WR. Augmentation of the Riboflavin-Biosynthetic Pathway Enhances Mucosa-Associated Invariant T (MAIT) Cell Activation and Diminishes Mycobacterium tuberculosis Virulence. mBio. 2021;13(1):e0386521.

17. Chengalroyen MD, Mehaffy C, Lucas M, Bauer N, Raphela ML, Oketade N, et al. Modulation of riboflavin biosynthesis and utilization in mycobacteria. Microbiol Spectr. 2024:e0320723.

18. Grinter R, Greening C. Cofactor F420: an expanded view of its distribution, biosynthesis and roles in bacteria and archaea. FEMS Microbiol Rev. 2021;45(5).

19. Macheroux P, Kappes B, Ealick SE. Flavogenomics--a genomic and structural view of flavin-dependent proteins. FEBS J. 2011;278(15):2625–34.

20. Selengut JD, Haft DH. Unexpected abundance of coenzyme F(420)-dependent enzymes in Mycobacterium tuberculosis and other actinobacteria. J Bacteriol. 2010;192(21):5788–98.

21. Kis K, Kugelbrey K, Bacher A. Biosynthesis of riboflavin. The reaction catalyzed by 6,7-dimethyl-8-ribityllumazine synthase can proceed without enzymatic catalysis under physiological conditions. J Org Chem. 2001;66(8):2555–9.

22. Lewinsohn DA, Winata E, Swarbrick GM, Tanner KE, Cook MS, Null MD, et al. Immunodominant tuberculosis CD8 antigens preferentially restricted by HLA-B. PLoS Pathog. 2007;3(9):1240–9.

23. Eckle SB, Corbett AJ, Keller AN, Chen Z, Godfrey DI, Liu L, et al. Recognition of Vitamin B Precursors and Byproducts by Mucosal Associated Invariant T Cells. J Biol Chem. 2015;290(51):30204–11.

24. Pethe K, Sequeira PC, Agarwalla S, Rhee K, Kuhen K, Phong WY, et al. A chemical genetic screen in Mycobacterium tuberculosis identifies carbon-source-dependent growth inhibitors devoid of in vivo efficacy. Nat Commun. 2010;1(5):57.

25. Rachman H, Kim N, Ulrichs T, Baumann S, Pradl L, Nasser Eddine A, et al. Critical role of methylglyoxal and AGE in mycobacteria-induced macrophage apoptosis and activation. PLoS One. 2006;1(1):e29.

26. Darwin KH, Ehrt S, Gutierrez-Ramos JC, Weich N, Nathan CF. The proteasome of Mycobacterium tuberculosis is required for resistance to nitric oxide. Science. 2003;302(5652):1963-6.

27. Narayanan GA, Nellore A, Tran J, Worley AH, Meermeier EW, Karamooz E, et al. Alternative splicing of MR1 regulates antigen presentation to MAIT cells. Sci Rep. 2020;10(1):15429.

28. Gopinath K, Warner DF, Mizrahi V. Targeted gene knockout and essentiality testing by homologous recombination. Methods Mol Biol. 2015;1285:131–49.

29. Murphy KC, Nelson SJ, Nambi S, Papavinasasundaram K, Baer CE, Sassetti CM. ORBIT: a New Paradigm for Genetic Engineering of Mycobacterial Chromosomes. mBio. 2018;9(6).

30. van Soolingen D, Hermans PW, de Haas PE, Soll DR, van Embden JD. Occurrence and stability of insertion sequences in Mycobacterium tuberculosis complex strains: evaluation of an insertion sequence-dependent DNA polymorphism as a tool in the epidemiology of tuberculosis. J Clin Microbiol. 1991;29(11):2578–86.

31. Mehaffy C, Lucas M, Kruh-Garcia NA, Dobos KM. Methods for Proteomic Analyses of Mycobacteria. In: Parish T, Kumar A, editors. Mycobacteria Protocols. New York, NY: Springer US; 2021. p. 533-48.

32. Yu F, Teo GC, Kong AT, Frohlich K, Li GX, Demichev V, et al. Analysis of DIA proteomics data using MSFragger-DIA and FragPipe computational platform. Nat Commun. 2023;14(1):4154.

33. MacLean B, Tomazela DM, Shulman N, Chambers M, Finney GL, Frewen B, et al. Skyline: an open source document editor for creating and analyzing targeted proteomics experiments. Bioinformatics. 2010;26(7):966-8.

34. Mehaffy C, Dobos KM, Nahid P, Kruh-Garcia NA. Second generation multiple reaction monitoring assays for enhanced detection of ultra-low abundance Mycobacterium tuberculosis peptides in human serum. Clin Proteomics. 2017;14:21.

35. Heinzel AS, Grotzke JE, Lines RA, Lewinsohn DA, McNabb AL, Streblow DN, et al. HLA-E-dependent presentation of Mtb-derived antigen to human CD8+ T cells. J Exp Med. 2002;196(11):1473–81.

